# *In vivo* host-pathogen interaction as revealed by global proteomic profiling of zebrafish larvae

**DOI:** 10.1101/140509

**Authors:** Francisco Díaz-Pascual, Javiera Ortíz-Severín, Macarena A. Varas, Miguel L. Allende, Francisco P. Chávez

## Abstract

The outcome of a host-pathogen interaction is determined by the conditions of the host, the pathogen, and the environment. Although numerous proteomic studies of *in vitro-grown* microbial pathogens have been performed, *in vivo* proteomic approaches are still rare. In addition, increasing evidence supports that *in vitro* studies inadequately reflect *in vivo* conditions. Choosing the proper host is essential to detect the expression of proteins from the pathogen *in* vivo. Numerous studies have demonstrated the suitability of zebrafish (*Danio rerio*) embryos as a model to *in vivo* studies of *Pseudomonas aeruginosa* infection. In most zebrafish-pathogen studies, infection is achieved by microinjection of bacteria into the larvae. However, few reports using static immersion of bacterial pathogens have been published. In this study we infected 3 days post-fertilization (DPF) zebrafish larvae with *P. aeruginosa* PAO1 by immersion and injection and tracked the *in vivo* immune response by the zebrafish. Additionally, by using non-isotopic (Q-exactive) metaproteomics we simultaneously evaluated the proteomic response of the pathogen (*P. aeruginosa* PAO1) and the host (zebrafish). We found some zebrafish metabolic pathways, such as hypoxia response via HIF activation pathway, exclusively enriched in the larvae exposed by static immersion. In contrast, we found that inflammation mediated by chemokine and cytokine signaling pathways was exclusively enriched in the larvae exposed by injection, while the integrin signaling pathway and angiogenesis were solely enriched in the larvae exposed by immersion. We also found important virulence factors from *P. aeruginosa* that were enriched only after exposure by injection, such as the Type-III secretion system and flagella-associated proteins. On the other hand, *P. aeruginosa* proteins involved in processes like biofilm formation, cellular responses to antibiotic and starvation were enriched exclusively after an exposure by immersion.

We demonstrated the suitability of zebrafish embryos as a model for *in vivo* host-pathogen based proteomic studies in *P. aeruginosa*. Our global proteomic profiling identifies novel molecular signatures that give systematic insight into zebrafish*-Pseudomonas* interaction.

## Introduction

*Pseudomonas aeruginosa* is one of the most common opportunistic pathogens in humans, normally infecting patients wounded, burned, immunocompromised or with cystic fibrosis. Many surrogate host models have been used to study the pathogenesis of *P. aeruginosa*, such as plants, amoebas, insects, fish and mice (Brannon et al., 2009; Mahajan-Miklos, Rahme, & Ausubel, 2000; Pukatzki, Kessin, & Mekalanos, 2002). Zebrafish (*Danio rerio*) combine the advantages of invertebrate and murine models. It has a similar immune system to that found in mammals, but without the costs and lab space requirements of murine models. It has a fully functional innate immune system in the first days of embryogenesis (Iwanami, 2014). However, the adaptive immune system is mature only after 4-6 weeks post-fertilization (Lam, Chua, Gong, Lam, & Sin, 2004).

The real-time visualization capabilities and genetic tractability of the zebrafish infection model allow the elucidation of molecular and cellular features of *P. aeruginosa* pathogenesis while causing damage to the host. Neutrophils and macrophages can rapidly phagocytize and kill *P. aeruginosa* (Herbomel, Thisse, & Thisse, 1999; Le Guyader et al., 2008), suggesting that both immune cell types play a role in the defense against bacterial infection.

There are two main methods for the exposure of the zebrafish to a bacterial pathogen, injection and static immersion. The former involves the injection of the bacteria directly into the fish and in the latter the zebrafish is incubated in a bacterial suspension. It is still unclear if the responses of zebrafish exposed to *P. aeruginosa* by these two methods are similar. Moreover, in contrast with early stages of development, there is little information regarding the response of 3 days post-fertilization (DPF) zebrafish larvae infected with this pathogen, age at which the innate immune system is already mature, the fish have left the corium and have opened their mouth.

A novel approach for host-pathogen studies on alternative models such as zebrafish infections is to study the whole spectrum of gene expression with techniques such as proteomics and transcriptomics. This enable us to recognize and analyze cellular processes from a global point of view and from these understand the molecular and cellular adjustments that occur during infection in both the pathogen and the host. However, limited studies have combined the *in vivo* cellular responses and global proteomic changes during host-pathogen interaction.

The aim of this study was to compare host-pathogen interaction during immersion and injection methods of exposure of zebrafish larvae to *P. aeruginosa* PAO1, and to identify marker genes for the immune response and virulence factors important for the infection. For this we have compared the *in vivo* neutrophil response and the global proteomic profiling of zebrafish exposed to *P. aeruginosa* PAO1 through injection in the caudal artery and through static immersion. By using Q-Exactive Orbitrap Mass Spectrometry, the global metaproteomic profiling of 3 DPF zebrafish larvae inoculated by both methods of infection was compared. The global proteomic profiling approach with zebrafish larvae infected with this pathogen allows for simultaneous tracking of the global proteome changes in the host (zebrafish) and in the bacterial pathogen (*P. aeruginosa*). We demonstrate that the global proteomic profiling we employed is a strong platform for *in vivo* global host-microbe interaction studies in zebrafish.

## Materials and Methods

### Zebrafish husbandry

Zebrafish (*Danio rerio*) embryos were obtained by natural spawning of Tab5 and *Tg(BACmpo:mCherry)* lines (Renshaw et al., 2006). Fertilized eggs were raised in petri dishes containing E3 medium (5 mM NaCl, 0.17 mM KCl, 0.33 mM CaCl_2_, 0.3 mM MgSO_4_) and 0.1% methylene blue until 72 hours post-fertilization (HPF). All procedures complied with national guidelines and were approved by the Animal Use Ethics Committee of the University of Chile and the Bioethics Advisory Committee of Fondecyt-Conicyt (the funding agency for this work).

### Bacterial immersion and injection experiments

*Pseudomonas aeruginosa* PAO1 was used for the injection and immersion assays. Avirulent *Escherichia coli* (strain DH5α) was used as a control. The bacteria were grown in LB medium overnight at 37°C and with shaking at 180 RPM. The cultures were washed and subsequently suspended in PGS medium (PGS (↓Pi); 2.5 g/L peptone, 3 g/L NaCl, 1 mM MgSO_4_, 1 mM CaCl_2_, 2% glycerol, pH 6.0); these suspensions were used to inoculate a liquid culture in PGS medium or PGS medium supplemented with inorganic phosphate (PGS (↑Pi); 25 mM potassium phosphate buffer pH 6.0) in a 1:100 ratio. These cultures were grown at 37°C with shaking at 180 RPM for 18 hrs.

For the immersion assays (Varas et al., 2017), cultures were washed and re-suspended in E3 medium (5 mM NaCl, 0.17 mM KCl, 0.33 mM CaCl_2_, 0.33 mM MgS0_4_, pH 7.0), then adjusted to an optical density at 600 nm (OD_600_) of 1.4 equivalent to (1.1 ± 0.23)E9 CFU/mL for PGS (↓Pi) and (1.1 ± 0.22)E9 CFU/mL for PGS (↑Pi) for *P. aeruginosa* and equivalent to (1.2 ± 0.25)E9 CFU/mL for PGS (↓Pi) and (1.0 ± 0.15)E9 CFU/mL for PGS (↑Pi) for *E. coli*. To determine CFU/mL, the bacterial suspensions were plated on LB-agar. Zebrafishes of 3 DPF were washed with sterile E3, and 10 larvae were placed per well in a 6 well plate. The wells were filled with the bacterial solution to obtain an adjusted OD_600_ of 0.7, 0.525 and 0.35 within a final volume of 8 ml, corresponding to (5.5± 0.22)E8 CFU/mL, (4.1± 0.20)E8 CFU/mL, (2.8± 0.23)E8 CFU/mL respectively. The zebrafish larvae were incubated at 20°C for 30 hrs.

For the injection assays, the bacterial cultures were washed and subsequently suspended in phosphate-buffered saline (PBS) and adjusted to an OD_600_ of 5.0. Zebrafish of 3 DPF were anesthetized with 0.01% tricaine and mounted in low melting point agarose. The zebrafish were kept under anesthesia for the duration of bacterial injections. Between 2,000 and 6,000 CFU were injected into the caudal artery. To determine the CFU injected, the droplets were also injected in sterile PBS and plated on LB-agar. The injected zebrafish were kept at 20°C for 72 hours.

Survival curves were analyzed using the Kaplan–Meier method (GraphPad Prism version 6.00 for Mac OS X, GraphPad Software, La Jolla California USA, www.graphpad.com). All experiments were done in triplicate for a total of 30 larvae per condition.

### Inflammation mediated by neutrophils

Recruitment of neutrophils was observed using the *Tg(BACmpo:mCherry)* zebrafish line. The larvae were observed using an Olympus MVX10 (Japan) fluorescence microscope at 2, 4, 7 and 22 hours post-injection (HPI). To determine the number of neutrophils in circulation, injected *Tg(BACmpo:mCherry)* zebrafish larvae were anesthetized with 0.01% tricaine for 2 minutes and the number of neutrophils that passed through the site of injection by the caudal artery in one minute were counted. Statistical differences were assessed using a two-way ANOVA followed by Turkey’s multiple comparisons test using GraphPad Prism version 6.00 for Mac OS X, GraphPad Software, La Jolla California USA, (www.graphpad.com).

### Global proteomic profiling using Q-exactive mas spectrometry

For the proteomic analysis we selected and froze samples in a dry methanol bath at -80°C, including: 10 larvae injected with 2,000-6,000 CFU of *P. aeruginosa* (grown in medium PGS (↓Pi)) at 22 HPI; 10 larvae injected with sterile PBS at 22 HPI; 10 larvae at 22 hours post-exposure (HPE) to ~2,5×10^8^ CFU/mL of *P. aeruginosa* (grown in medium PGS (↑Pi)) by static immersion; and 10 larvae incubated for 22 hours in sterile E3 medium.

Global proteomic profiles were obtained by services from Bioproximity, LLC (USA). Protein denaturation, digestion and desalting of samples were prepared using the filter-assisted sample preparation (FASP) method (Wiśniewski, Zougman, Nagaraj, & Mann, 2009). Briefly, the samples were digested using trypsin, and each digestion mixture was analyzed by UHPLC-MS/MS and a quadrupole-Orbitrap mass spectrometer (Q-Exactive, Thermo Fisher). Mass spectrometer RAW data files were compared with the most recent protein sequence libraries available from UniProtKB. Proteins were required to have 1 or more unique peptides across the analyzed samples with E-value scores of 0.0001 or less.

The differences between the proteome of the larvae injected with *P. aeruginosa* and PBS, and between the larvae exposed to *P. aeruginosa* by static immersion were determined. Significance cut-offs for the ratios were set at 1.5-fold change. The proteins up-regulated and down-regulated exclusively in one of the conditions were determined. Gene ontology (GO) analysis were performed using Panther web-based software (Mi, Muruganujan, Casagrande, & Thomas, 2013; Mi, Muruganujan, & Thomas, 2013). For enrichment analysis the cut of was set to P-value<0.05.

To analyze the *P. aeruginosa* proteome, proteins in the infected larvae but not in the respective control were analyzed using Panther web-based software and a cut-off set to P-value<0.05. For visualizing changes in the GO categories in both methods, Treemap (Version 3.7.2 for Mac) were used for the hierarchical representation of enriched biological processes and cellular components. Items were grouped by category and their size was proportional to the P-value.

## Results

### Neutrophil response toward *P. aeruginosa* PAO1 in zebrafish larvae after inoculation by microinjection and static immersion

*P. aeruginosa* becomes more virulent when grown with limited inorganic phosphate (Pi) in the medium (Long, Zaborina, Holbrook, Zaborin, & Alverdy, 2008; Zaborin et al., 2012). Therefore, in order to obtain a wide range of outcomes to compare the injection and static immersion methods, the bacterial cells were grown in PGS medium lacking Pi or in the same medium supplemented with Pi, PGS (↓Pi) or PGS (↑Pi) respectively.

For the immersion assay, zebrafish larvae at 72 HPF were immersed in high concentrations of *P. aeruginosa* or *E. coli* (in the order of 10^8^ CFU/ml) for 30 hours at 20°C. Mortality of zebrafish was observed when they were immersed with *P. aeruginosa*, but not with *E. coli* (Figures 1A and 1B). As expected, the larvae survival was dependent on the concentration of bacteria and the virulence of *P. aeruginosa* was increased when they were grown in Pi limitation. Before the death of the larvae, their tail was curved and there was damage and necrosis in the fins, tail and head (Figure 1C). Using the transgenic line *Tg(BACmpo:mCherry)* we did not find significant migration of neutrophils or changes in the distribution of these cells when the zebrafish was immersed with *P. aeruginosa* (Figure 1D).

**Figure 1.**
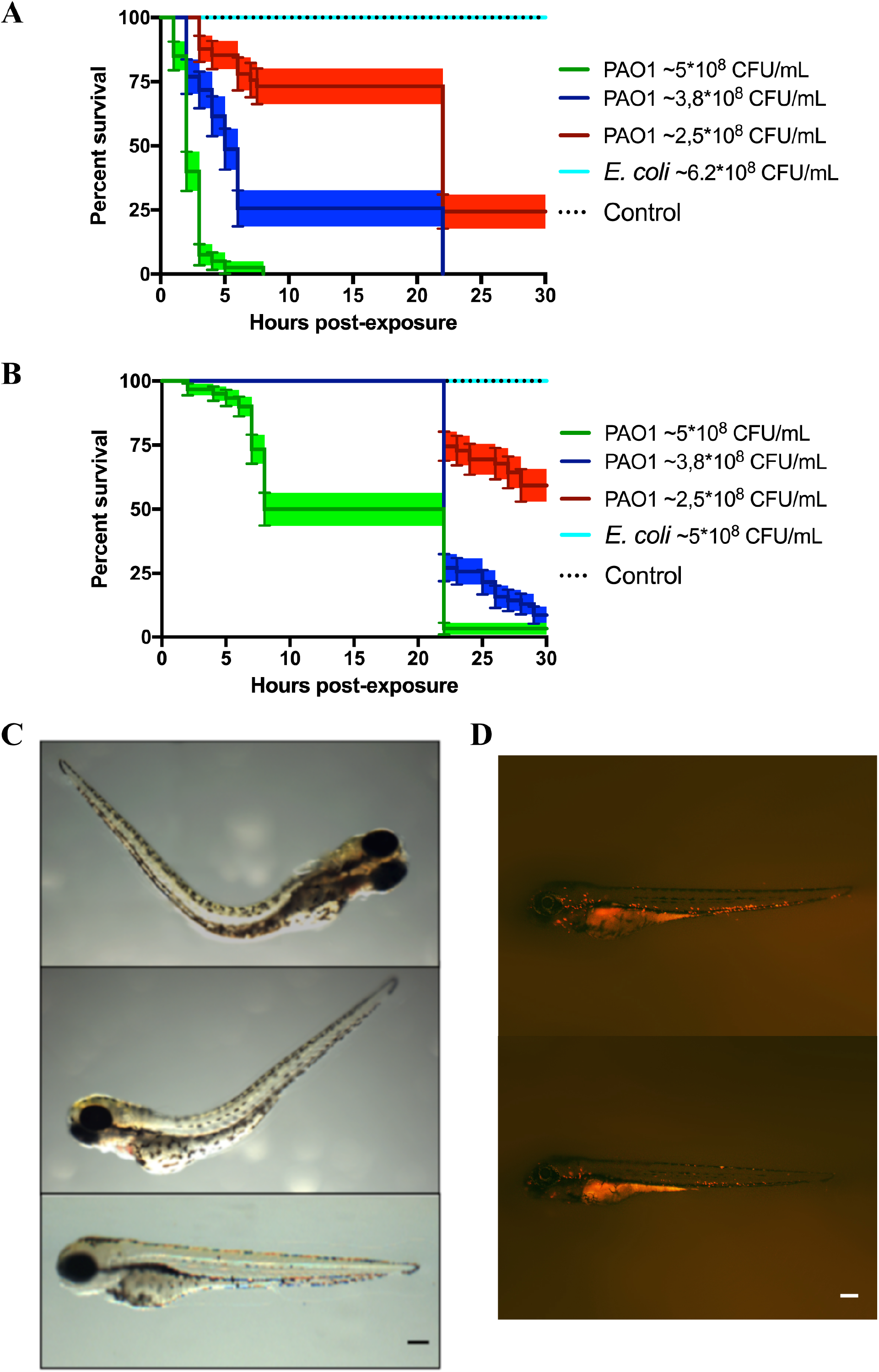
Zebrafish larvae exposed to *P. aeruginosa* by static immersion. Larvae were immersed at 72 HPF in *P. aeruginosa* PAO1, *E. coli* DH5α or in sterile E3 medium as control. (A, B) Survival curve of 72 HPF larvae immersed in different suspensions *P. aeruginosa* PAO1 (green, blue and red) or in a suspension of *E. coli* DH5α (light blue line) or in sterile E3 medium (black dotted line). In (A) the bacteria were grown in PGS (↓Pi) medium. In (B) the bacteria were grown in PGS (↑Pi) medium. (C) Larvae immersed with ~5x10^8^ CFU/mL of *P. aeruginosa* PAO1 grown in PGS (↓Pi) medium (upper and second picture) or in sterile E3 medium (bottom picture) at 3 hpe. (D) *Tg(BACmpo:mCherry)* larvae immersed at 72 HPF in ~2,5x10^8^ CFC/mL of *P. aeruginosa* PA01 grown in (↑Pi) medium (left) or in sterile E3 medium (right) at 22 hpe. Scale A-B: 100 μm.

The region of the caudal artery was chosen as the site of injection due to the reduced pigmentation in comparison to the caudal vein. Between 2,000-6,000 CFU of *P. aeruginosa* or *E. coli* cells were injected in zebrafish larvae at 72 HPF, and sterile PBS was injected as control. Under the tested conditions, death of the larvae was observed only when *P. aeruginosa* grown in PGS (↓Pi) was injected (Figure 2A). Several larvae injected with *P. aeruginosa* presented damage and necrosis in the tail before their death (Figure 2B). In the first HPI, a circulatory blockage at the site of infection was observed when injected with *P. aeruginosa*, *E. coli* or sterile PBS. However, this blockage remained only for a few hours in the larvae injected with PBS sterile, but it was observable until 28 HPI in the larvae injected with *P. aeruginosa*.

**Figure 2.**
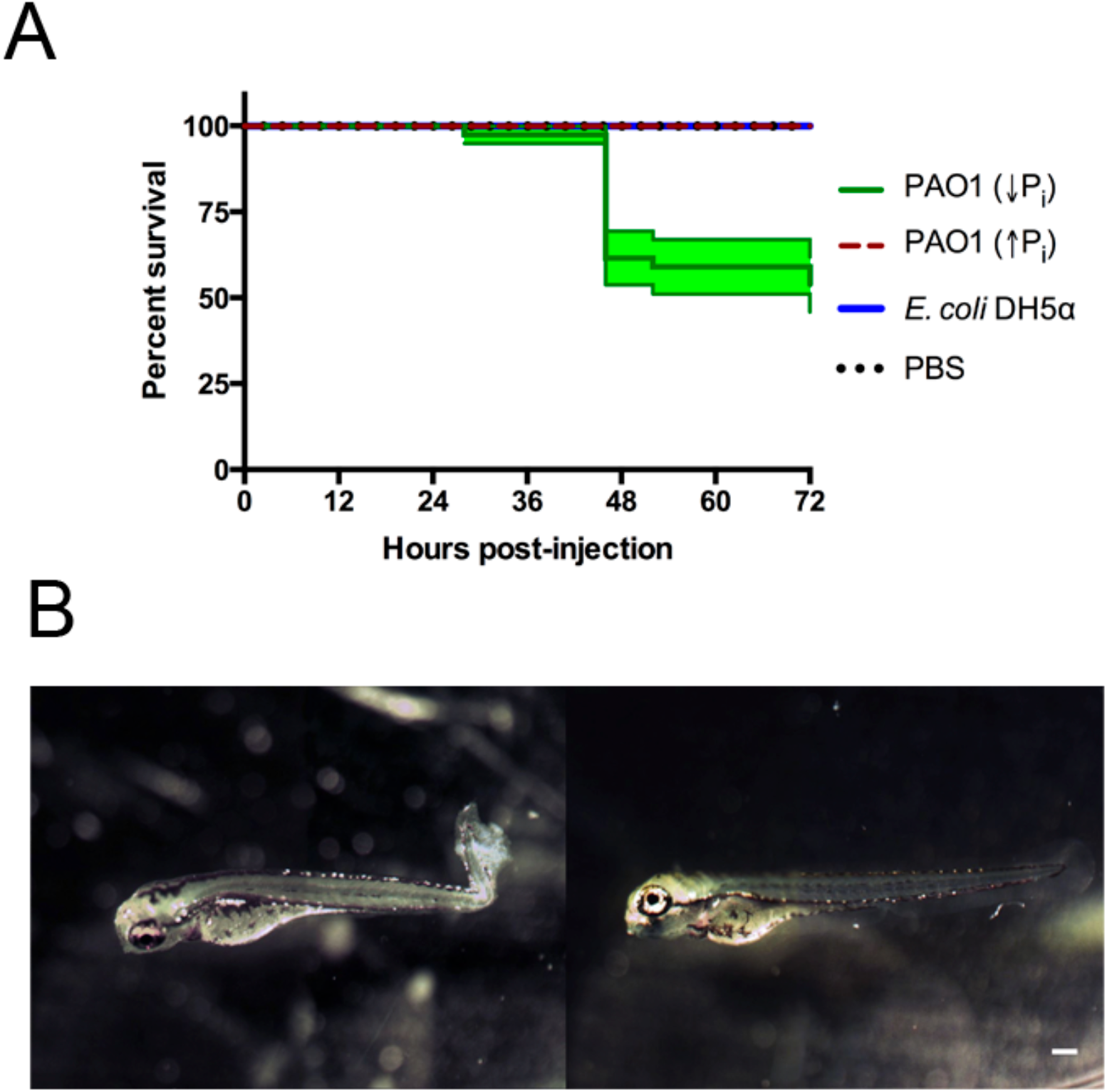
Zebrafish larvae exposed to *P. aeruginosa* by injection. Larvae were injected at 72 HPF with 2,000-6,000 CFU of *P. aeruginosa* PAO1 or *E. coli* DH5α into the caudal artery. Sterile PBS was injected as control. (A) Survival curve of 3 DPF larvae injected with *P. aeruginosa* PAO1 grown in PGS (↓Pi) medium (green line), *P. aeruginosa* PAO1 grown in PGS (↑Pi) medium (red dashed line), *E coli* DH5α (blue line) or sterile PBS medium (black dotted line). (B) Larvae injected at 72 HPF with *P. aeruginosa* PAO1 grown in PGS (↓Pi) medium (left) or injected with sterile PBS medium (right) at 28 HPI. Scale: 100 μm.

In order to compare both methods of exposure and considering the absence of zebrafish mortality before 40 HPI, we address if in previous HPI there was some response in the larvae, specifically an inflammatory response. To do so, a transgenic line that has the neutrophils marked with the fluorescent protein mCherry was used and injected with *P. aeruginosa*, *E. coli* or sterile PBS. In the larvae injected with *P. aeruginosa* there was a clear recruitment of neutrophils to the site on injection that remained until 28 HPI. This also occurred after an injection with sterile PBS, however under these conditions it was barely observable at 6 HPI and it was totally absent at 28 HPI (Figure 3B). These results suggest that an injection with *P. aeruginosa* generates an important inflammatory response at the site of injection. To quantify this, the number of neutrophils that passed trough the site of injection in one minute was determined (Figure 3A). At 22 HPI and 28 HPI there was a significant difference between the number of neutrophils circulating in the larvae injected with *P. aeruginosa* grown in PGS (↓Pi) or injected with *E. coli* with respect to the control. In contrast, there was no difference between the larvae injected with *P. aeruginosa* grown in PGS (↑Pi) with those injected with *E. coli*.

**Figure 3.**
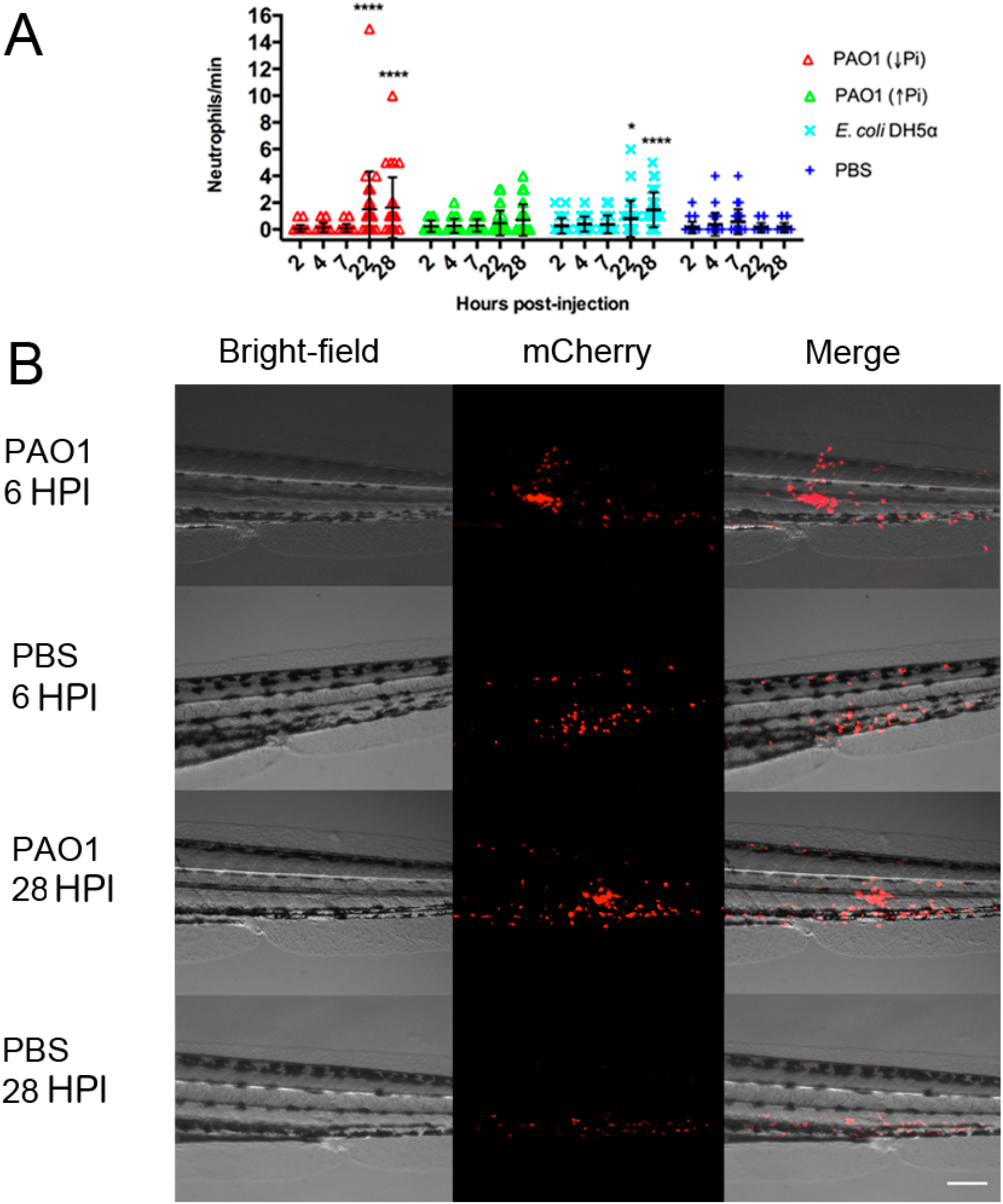
Inflammation mediated by neutrophils in injected zebrafish. *Tg(BACmpo:mCherry)* larvae were injected at 72 HPF with 2,000-6,000 CFU of *P. aeruginosa* PAO1 or *E. coli* DH5α into the caudal artery. Sterile PBS was injected as control. (A) Larvae were injected with *P. aeruginosa* PAO1 grown in PGS (↓Pi) medium (red), *P. aeruginosa* PAO1 grown in PGS (↑Pi) medium (green), *E coli* DH5α (light blue) or sterile PBS medium (blue). Neutrophils that passed through the site of injection by the caudal artery in one minute at 2, 4, 7, 22 and 28 HPI were counted. Each symbol represents a different zebrafish larva. Statistical differences with the control were determined. * P≤0.05. **** P≤0.0001. (B) *Tg(BACmpo:mCherry)* larvae injected at 72 hpf with *P. aeruginosa* grown in PGS (↓Pi) medium or sterile PBS medium were imaged at 6 and 28 HPI at the site of injection. Scale: 100 μm

### Global proteomic profiling after infection by microinjection and static immersion

First, the zebrafish proteomes were analyzed. The proteome profile of the larvae injected with *P. aeruginosa* was compared with larvae injected with sterile PBS (Figure 4A). A total of 12,677 proteins were up-regulated (fold-change>1.5) and 13,292 proteins were significantly down-regulated (fold change <-1.5). From these, 8,575 proteins were up-regulated and 9,363 proteins were down-regulated exclusively in this condition, and therefore not in the larvae exposed by immersion. On the other hand, when the proteome of the larvae immersed with *P. aeruginosa* was compared to the proteome of the larvae immersed in sterile E3 medium (control), a total of 13,607 proteins were up-regulated and 13,136 proteins were down-regulated. From these 9,505 proteins were up-regulated and 9,208 proteins were down-regulated exclusively in the larvae exposed by immersion.

**Figure 4.**
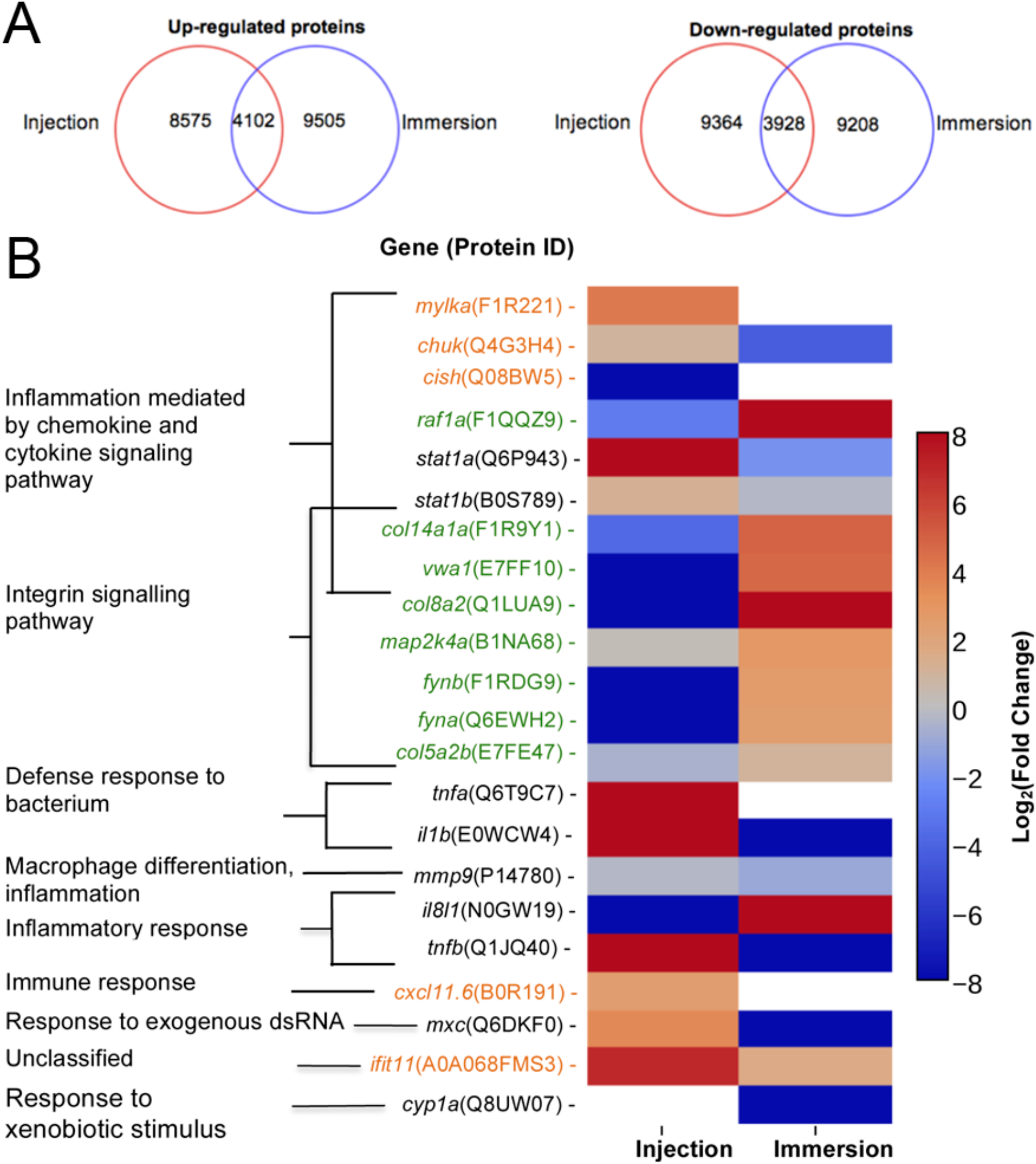
Proteins up-regulated and down regulated according to method of exposure and molecular markers proposed. (A) Number of proteins up-regulated (right) or down-regulated (left) in zebrafish exposed at 72 HPF to *P. aeruginosa* PAO1 by injection (inside red circle) or static immersion (inside blue circle) at 22 hpe. (B) Molecular markers proposed for a *P. aeruginosa* PAO1 infection by injection (orange) or by static immersion (green) in 3 DPF zebrafish. In black there are the markers previously reported for other bacterial pathogens. The category displayed for the markers was determined using the homologous gene in human.

The proteins up-regulated and down regulated in each condition were categorized according to GO-terms of biological process (Supplementary Files 1-3). No appreciable differences between the proteins up-regulated and down-regulated were found in each condition and neither between conditions in regard of biological process. The majority of the proteins were categorized under “metabolic process (GO:0008152)” and cellular process (GO:0009987)” with approximately 25% and 20% respectively in all the analysis. The GO-term “immune system process (GO:0002376)” represented only around 5% of the proteins and the GO-term “response to stimulus” only around 6% of the proteins.

Afterwards, an overrepresentation test was carried out and the proteins were analyzed by Pathway (Table 1). In the up-regulated proteins of the larvae exposed by static immersion the “hypoxia response via HIF activation pathway (P00030)” was enriched. In contrast, in the down-regulated proteins of the larvae exposed by injection the categories “B cell activation (P00010)” and “endothelin signaling pathway (P00019)” were enriched. There were no significantly enriched groups in the up-regulated proteins by injection.

**Table 1.** Overrepresentation test of proteins up-regulated or down-regulated in larvae exposed to *P. aeruginosa* according to Pathway. Up-regulated and down-regulated proteins in zebrafish injected with *P. aeruginosa* PAO1 or co-incubated with *P. aeruginosa* PAO1, with respect to the controls, were analyzed according to Pathway.

| Pathway | Proteins found | Expected proteins | Fold enrichment | P-value |
| --- | --- | --- | --- | --- |
| **Proteins up-regulated in zebrafish exposed by immersion** |  |  |  |  |
| **Hypoxia response via HIF activation** | 21 | 8.15 | 2.58 | 1.90E-02 |
| **Proteins down-regulated in zebrafish exposed by immersion** |  |  |  |  |
| **Integrin signaling pathway** | 64 | 40.03 | 1.6 | 4.41E-02 |
| **Proteins down-regulated in injected zebrafish** |  |  |  |  |
| **B cell activation** | 34 | 16.34 | 2.08 | 1.38E-02 |
| **Endothelin signaling pathway** | 41 | 22.61 | 1.81 | 4.96E-02 |
| **Proteins up-regulated only in injected zebrafish** |  |  |  |  |
| **Axon guidance mediated by netrin** | 21 | 6.95 | 3.02 | 2.03E-03 |
| **Cytoskeletal regulation by Rho GTPase** | 32 | 16.13 | 1.98 | 4.83E-02 |
| **Inflammation mediated by chemokine and cytokine signaling pathway** | 58 | 34.34 | 1.69 | 2.11E-02 |
| **Proteins up-regulated only in zebrafish exposed by immersion** |  |  |  |  |
| **Integrin signaling pathway** | 50 | 28.91 | 1.73 | 3.49E-02 |
| **Angiogenesis** | 53 | 31.28 | 1.69 | 3.73E-02 |

To address the differences in the proteins that changed their levels in one method of exposure but not in the other, the proteins up-regulated only in larvae exposed by injection or in larvae exposed by immersion were analyzed by Pathway. In the proteins up-regulated exclusively in injected larvae the pathways “axon guidance mediated by netrin (P00009)”, “inflammation mediated by chemokine and cytokine signaling pathway (P00031)” and “Cytoskeletal regulation by Rho GTPase (P00016)” were enriched. On the other hand, in the proteins up-regulated exclusively in the larvae exposed by immersion, “integrin signaling pathway” and “angiogenesis (P00005)” were enriched. No pathways were significantly enriched when proteins that were down-regulated exclusively in one of the conditions or the proteins that were up-regulated in both conditions were analyzed.

Several genes or gene products that have been reported as genetic markers for exposure to bacteria by injection were found in the zebrafish proteome (Figure 4B). Particularly, the levels of the transcripts of the *mmp9* and *il8l1*genes were reported as up-regulated in zebrafish after an exposure to *S. typhimurium* by injection (Stockhammer, Zakrzewska, Hegedus, Spaink, & Meijer, 2009). However, neither of the proteins encoded by these genes were significantly up-regulated in larvae injected with *P. aeruginosa* at 22 HPI. IL-8L1 (N0GW19) was only detected in the larvae immersed with *P. aeruginosa* and in the larvae injected with PBS, but not in the control immersed in E3 medium or injected with *P. aeruginosa*. MMP9 (P14780) was slightly down-regulated (−2.04-fold change) in the larvae exposed by static immersion. The proteins IL-1ß (E0WCW4), TNF-ß (Q1JQ40), interferon-induced GTP-binding protein MxC (Q6DKF0), STAT1A and STAT1B were reported as molecular markers in zebrafish injected either with *S. Typhimurium* (Stockhammer et al., 2009) or *E. tarda* (van Soest et al., 2011). We found all these proteins up-regulated in the larvae injected with *P. aeruginosa*. Interestingly, they were all down-regulated in the larvae immersed in *P. aeruginosa*, except for STAT1B (−1.24-fold change). In addition, IL-1ß and TNF-α were reported as molecular markers for the response of the zebrafish injected at 50 HPF with *P. aeruginosa* PA14 into the yolk circulation valley (Clatworthy et al., 2009). Both proteins were up-regulated in our injection assay, but not in the larvae exposed by static immersion. These results suggest that the expression of known components in the response towards a bacterial injection is dependent on the types of pathogens infected. To our knowledge, the expression of *cyp1a* is the only proposed marker for an exposure by immersion to a *E. tarda* (van Soest et al., 2011). However, in this study CYP1A was only detected in the larvae incubated in sterile E3 medium.

To suggest potential novel molecular markers that are exclusive either for the injection or static immersion methods for exposure we look for proteins that fulfill the following conditions: (1) the protein should participate in pathways that are solely enriched in the larvae infected by one of the exposure methods, (2) the protein or its human homologue should have a known biological function related with an infective process and (3) the levels of protein expression be distinct in each method of exposure.

Considering these conditions, we suggest that proteins MYLKA, CHUK, CXCL11.6, CISH and IFIT5 can be used as noel molecular markers for the injection method (Figure 4B). MYLKA had a 16.7-fold change were it was compared to the control. CHUK was up-regulated in injected larvae (1.93-fold change). CXCL11.6 had a 5.30-fold change in injected larvae. IFIT5 had a 121-fold change. The proteins COL8A2, COL141A, COL5A2B, VWA1, FYNA, FYNB, RAF1A and MAP2KA are potential novel markers for an infection by static immersion (Figure 4B). These proteins were significantly up-regulated in the larvae exposed by static immersion and down-regulated in the injected larvae, except for MAP2KA that was not down-regulated in the injected larvae (1.14-fold change).

We next analyzed *P. aeruginosa* proteomes to address the differences in the proteins expressed in the *P. aeruginosa* injected and the *P. aeruginosa* in the immersion assay. A total of 1,159 *P. aeruginosa* PAO1 proteins were exclusively found in the injected larvae, and 1,276 in the larvae exposed by immersion (Supplementary Table 2). All of these proteins were analyzed by the GO-terms of component and biological process (Figure 5, Supplementary Table 3). Several GO-term categories associated with the bacterial-type flagellum were enriched in the *P. aeruginosa* injected, but not in the immersion assay. In regard of the GO-term by biological process, the categories “protein secretion by the type III secretion system” (GO:0030254), “bacterial-type flagellum organization” (GO:0044781), “putrescine metabolic process” (GO:0009445) and “response to drug” (GO:0042493) were enriched in the injected *P. aeruginosa*, but not in the immersion assay. On the other hand, the categories “single-species biofilm formation” (GO:0044010), “pathogenesis” (GO:0009405), “cellular response to antibiotic” (GO:0009267) and “cellular response to starvation” (GO:0009267) were enriched in the *P. aeruginosa* co-incubated with the zebrafish, but not in the injected bacteria. The categories “biological adhesion” (GO:0022610), “cellular response to stress” (GO:0033554), “pilus assembly” (GO:0009297), “biofilm formation” (GO:0042710), “cell communication” (GO:0007154), “type IV pilus-dependent motility” (GO:0043107), “cilium or flagellum-dependent cell motility” (GO:0001539), “protein secretion by the type II secretion system” (GO:0015628), “secondary metabolite biosynthetic process” (GO:0044550), “siderophore metabolic process” (GO:0009237), “pyoverdine metabolic process” (GO:0002048), “response to antibiotic” (GO:0046677) and “peptidoglycan metabolic process” (GO:0000270) were enriched in both groups of proteins (Supplementary Table 3).

**Figure 5.**
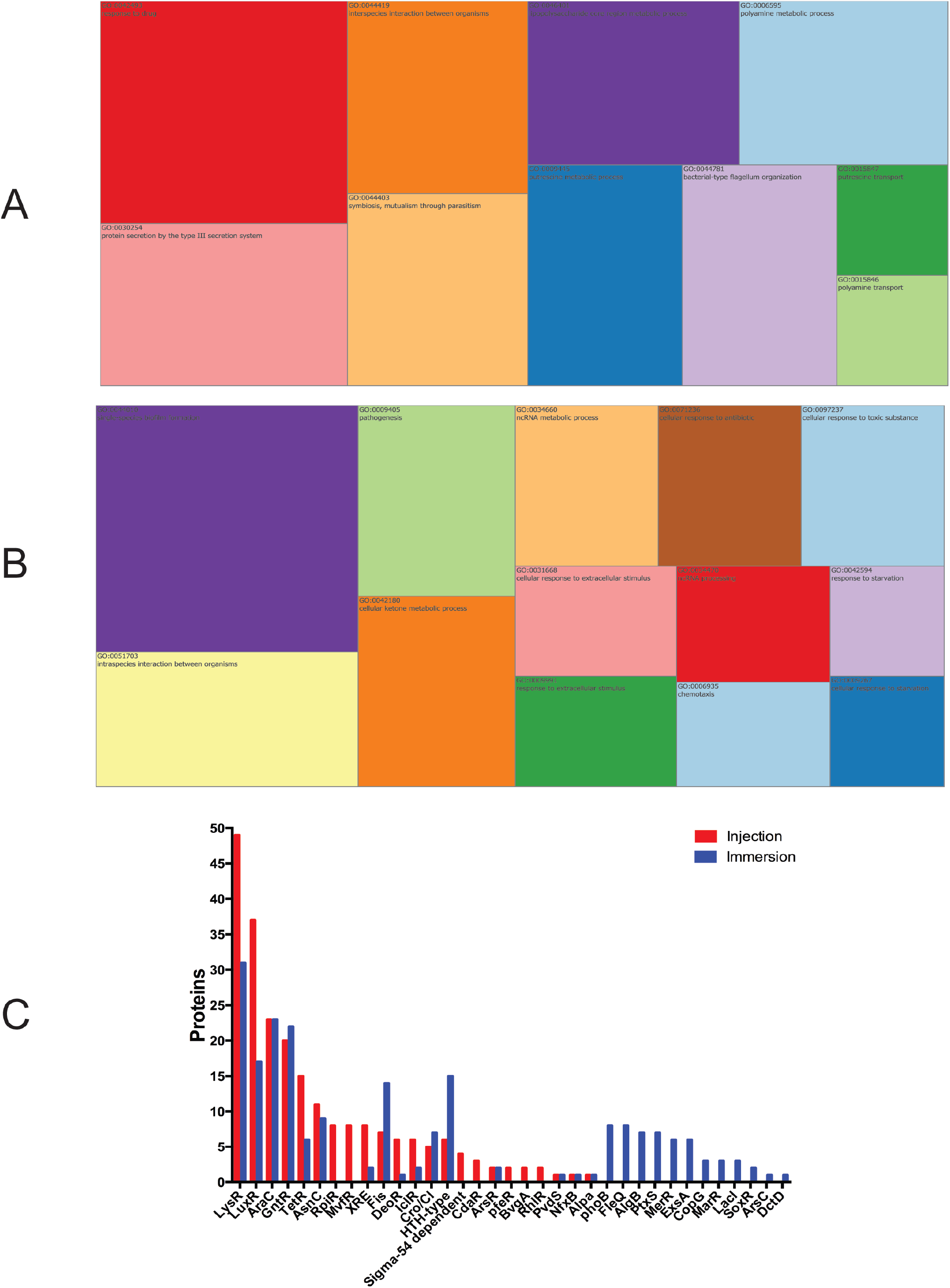
*P. aeruginosa* proteins exclusively found in zebrafish infected by injection (A) or by static immersion. Treemap of the overrepresentation analysis of *P. aeruginosa* proteins exclusively found in zebrafish infected by injection (A) or by static immersion (B). (C) Representation of transcription factors detected among these exclusively found proteins in each method. The size of each group is proportional to the P-value. List of proteins were categorized by biological process (GO). The cut-off was set at P<0.05.

To explain how these processes were regulated after an infection by injection or static immersion, the bacterial transcription regulators detected in the infected larvae but not in the respective controls were analyzed (Figure 5B). A total of 15 bacterial transcription regulator families were detected in the larvae injected with *P. aeruginosa* and in the larvae exposed by static immersion (LysR, LuxR, AraC, GntR, TetR, AsnC, XRE, Fis, DeoR, IclR, Cro/CI, HTH-type, ArsR, PvdS and AlpA), which suggest that several processes are regulated in a similar way in both methods of exposure. A total of 7 bacterial transcription factor families were detected only in the in the larvae injected with *P. aeruginosa* (RpiR, MvfR, Sigma-54 dependent family, PfeR, BvgA and RhlR). 12 bacterial transcription regulator families were detected only in the larvae in the larvae exposed by static immersion (PhoB, FleQ, AlgB, PtxS, MerR, ExsA, CopG, MarR, LacI, SoxR, ArsC, and DctD).

## Discussion

Zebrafish has received increasingly strong support as a surrogate to murine models for infection research (Lieschke & Currie, 2007; Tobin, May, & Wheeler, 2012). To enable high-throughput screening, a simple infection method will provide numerous benefits. In this report we have compared the neutrophil response and global proteome profile of zebrafish larvae infected by static immersion in *P. aeruginosa* with the profiles of fish into which bacteria were directly injected.

The injection of bacteria into the zebrafish generated an inflammatory response, which was absent in larvae exposed by immersion. This was observed by an increase in the neutrophils in circulation and a strong neutrophil recruitment to the site of injection. This response was similar after an injection with *E. coli* or *P. aeruginosa* grown in PGS (↓Pi). Although *E. coli* is non-virulent, it produces LPS, which causes an inflammatory response in the zebrafish upon exposure (Watzke, Schirmer, & Scholz, 2007). These results suggest that the zebrafish generates an immune response towards a bacterial injection by increasing the level of neutrophils in circulation. The injected larvae with *P. aeruginosa* grown in PGS (↓Pi) were damaged, presenting necrosis in their tails. This effect has been already reported after injections of *P. aeruginosa* PAO1 into the caudal vein of zebrafish at 28 HPF (Llamas et al., 2009), which suggests that this phenotype is not exclusive for this site of infection or the age of the larvae fish.

All injected larvae also presented a circulatory blockage. However, this phenotype lasted longer if the injection was with *P. aeruginosa*. This phenomenon is not exclusive for an injection into the caudal artery, as it has been also observed when *P. aeruginosa* PAO1 was injected into the caudal vein of zebrafish embryos at 50 HPF; in this previously published case, circulatory blockage was caused by an aggregation of blood cells and bacteria (Brannon et al., 2009). Among the down-regulated proteins of the larvae exposed by injection, the B cell activation and endothelin signaling pathways were enriched. At this stage of development, the zebrafish does not have a functional adaptive immune system; therefore a decrease in the proteins related to the activation of B cell may be due to a regulation process in the precursors of this cell lineage. The endothelin signaling pathway is involved with blood pressure regulation, specifically by contracting the blood vessels (Marteau, Zaiou, Siest, & Visvikis-Siest, 2005). A decrease of the proteins involved in this pathway could be a way to diminish the blood pressure produced by the damage and circulatory blockage generated by the bacterial injection.

On the other hand, in the larvae exposed by the immersion method, the hypoxia response via HIF activation pathway was enriched, which suggests that the larvae suffered from a lack of oxygen when exposed to *P. aeruginosa* by static immersion, possible caused by consumption of oxygen by the bacterial population and generation of compounds produced that diminish the percentage of dissolved oxygen, such as cyanide. Metabolic pathways related to angiogenesis were enriched, which could be due to the well-documented relation between hypoxia response and angiogenesis (Krock, Skuli, & Simon, 2011). Among down-regulated proteins, the integrin signaling pathway was enriched, which could be related to cell migration, especially to the epithelia. This pathway is involved in cell attachment, and down-regulation of this process could be a response against bacteria that are outside the fish or in contact with its skin.

Among the several host markers previously reported in zebrafish (Clatworthy et al., 2009; Stockhammer et al., 2009; van Soest et al., 2011), only IL-1ß and TNF-α were found up-regulated in this study. This suggests that the zebrafish response depends on the host-pathogen interaction (e.g. age of the fish and pathogen). CYPA1, the only known marker for an exposure by immersion was not detected in the infected larvae. This could be explained since the expression of *cyp1a* had been reported as transient and mainly induced in the first hours of the exposure (van Soest et al., 2011).

For the injection method we suggest the proteins MYLKA, CHUK, CXCL11.6, CISH and IFIT5 as novel molecular markers for *P. aeruginosa* infection in zebrafish (Figure 4B). The human homologue of MYLKA is a calcium/calmodulin-dependent myosin light chain kinase, which is involved in the inflammatory response, and variants of this proteins increase the risk of acute lung injury (Gao et al., 2006). CHUK is involved in the macrophage inflammatory response toward LPS peritonitis in mouse (Kanaan et al., 2012), and it is important for functional maturation of dendritic cells (Mancino et al., 2013). CXCL11.6 is homologue of CXCL11 in humans, which is involved in the chemotaxis of interleukin activated T cells. In zebrafish this interleukin is involved in the macrophage recruitment after a mycobacteria infection (Torraca et al., 2015). The CISH homologue in humans is involved in the negative regulation of cytokines. This protein was not detected in the larvae injected with *P. aeruginosa*, but it was detected in the larvae injected with PBS medium. This observation suggests that its diminished levels could be important for the zebrafish immune response. The expression of the gene *cish* was also diminished in catfish after bacterial infection (Yao et al., 2015). IFIT11, the zebrafish homologue of human IFIT15, is involved in the response toward an intramuscular viral injection in salmons (Chang, Robertsen, Sun, & Robertsen, 2014). Interestingly, its levels had a 121-fold change in the larvae injected with *P. aeruginosa*, which suggests that it also has a role towards a bacterial infection.

On the other hand, the proteins COL8A2, COL141A, COL5A2B, VWA1, FYNA, FYNB, RAF1A and MAP2KA could act as novel markers for an infection by static immersion. The genes *col8a2, col14a1a* and *col5a2b* are homologues of genes that encode for collagen in mice and humans. Since after an exposure by static immersion there was damage of the skin and external membranes, these proteins could be involved in the zebrafish defense towards that kind of damage. COL8A2 is involved in cellular migration and integrity of blood vessels. Morpholino knock-down of COL14A1A provokes skin detachment in mice (Bader et al., 2013), and therefore is thought to be involved in maintaining the integrity of skin and membranes. Zebrafish *col5a2b* is homologue of *col5a2* in mice, which is involved in wound healing and in the *in vitro* response of macrophage toward *Porphyromonas gingivalis* infection (Richard et al., 2013). The human homologue of the gene *vwa1* is hypothesized to be involved in extracellular matrix organization and behavioral response to pain. MAP2K4, the human homologue of MAP2K4A is required for maintaining peripheral lymphoid homeostasis and is involved in human gingival fibroblasts’ *in vitro* response to *P. gingivalis* exposure (Herath et al., 2013). The gene *fyn*, a homologue of *fyna* and *fynb*, encodes a kinase involved in the control of cell growth, axon guidance, signaling by integrin, immune response and cellular migration. In mice *fyn* is expressed in T cells and is involved in T cell receptor signaling and production of interleukin 4 (Mamchak et al., 2008). RAF1, the human homologue of the protein encoded by *raf1a* in zebrafish, is involved in signaling pathways that regulate apoptosis, angiogenesis and cellular migration. RAF1A levels diminished when the zebrafish are challenged with a viral infection (Encinas et al., 2013).

In regard of the bacterial proteome, the LysR family was the family with more transcription regulator found in both group of larvae (41 proteins in the injected larvae and 31 proteins in the immersed larvae), which is expected since the LysR-type Regulator Transcriptional Regulators (LTTRs) are one of the largest and most abundant classes of transcription regulator in prokaryotes. They are involved in the regulation of motility, metabolism, quorum sensing and virulence (Maddocks & Oyston, 2008). MvfR is a LTTR and was only detected in the injected larvae (8 different proteins ID). It is involved pathogenesis by the biosynthesis the pyocyanin, elastase, phospholipase and quorum-sensing related molecules (Cao et al., 2001). AlgB is involved in the production of alginate and is necessary for the high production of this exopolysaccharide in mucoid strains (Wozniak & Ohman, 1991). In these proteomic results, AlgB was only detected in the larvae exposed by immersion. The production alginates protects the bacteria form antibiotics and from the host immune response (Leid et al., 2005).

Under the conditions tested in this study, the injection method is more suitable to study the response of the host towards a bacterial infection, because it generates an immune response by inflammation and neutrophil recruitment that do not fish subjected to the immersion method. In contrast, the latter method is preferable to study the bacterial pathogenesis due to simplicity and the easiness to modify the bacterial conditions, which is desired to study bacterial virulence.

Finally, our global proteomic approach allows for systematic searches for the virulence factors that *P. aeruginosa* expresses *in vivo* during the infection process. Several known virulence factors from *P. aeruginosa* PAO1 (Pukatzki et al., 2002) toxin production, type II secretion, quorum sensing, production of extracellular polymeric substances – among many others – were identified *in vivo* in our proteomic studies. This highlights the importance of using surrogate host models for large-scale infection and immunization studies that can be further investigated in murine models (Kurz and Ewbank, 2007).

In summary, our global proteomic profiling study of injected larvae indicates an inflammatory response, which does not occur among larvae subjected to immersion in medium colonized by the same pathogenic bacteria. Our data suggest that the immersion method may cause an epithelial or other tissue response towards molecules that are shed or secreted by *P. aeruginosa*. Particularly interesting is that the expression of several toxic pigments including siderophores was augmented exclusively in the infected zebrafish. Therefore, combining live cell imaging with global proteomic profiling analysis in zebrafish larvae will be useful for future analysis of signal transduction pathways underlying host-pathogen interaction.

## Author contributions

Conceived and designed the experiments: FDP and FC. Performed the experiments: FDP, MV and JOS. Analyzed the data: FDP, MV, JOS, MA and JOS. Contributed with reagents/animals/materials/analysis tools: FC and MA. Wrote the paper: FDP and FC. All authors read and approved the final manuscript.

## Acknowledgments

We are indebted to Nicole Molina for technical support at SysmicroLab.

## Supplementary Material

**Supplementary Table 1. Overrepresentation test of proteins up-regulated and down-regulated in larvae exposed to *P. aeruginosa* according to Biological process**. The up-regulated and down-regulated proteins in zebrafish injected with *P. aeruginosa* PAO1 or co-incubated with *P. aeruginosa* PAO1, with respect to the controls, were analyzed according to Biological Process. The cut of was set to P-value<0.05

**Supplementary Table 2. Lists of *P. aeruginosa* proteins exclusively found in zebrafish infected by injection (Tab 1) or by static immersion (Tab 3)**. List of transcription factors detected among these proteins (Tab 3 corresponds to injection Tab4 to immersion). The proteins are accompanied by their gene name, description, average intensity, expected value, tax id from the NCBI database and KEGG database.

**Supplementary Figure 1.**
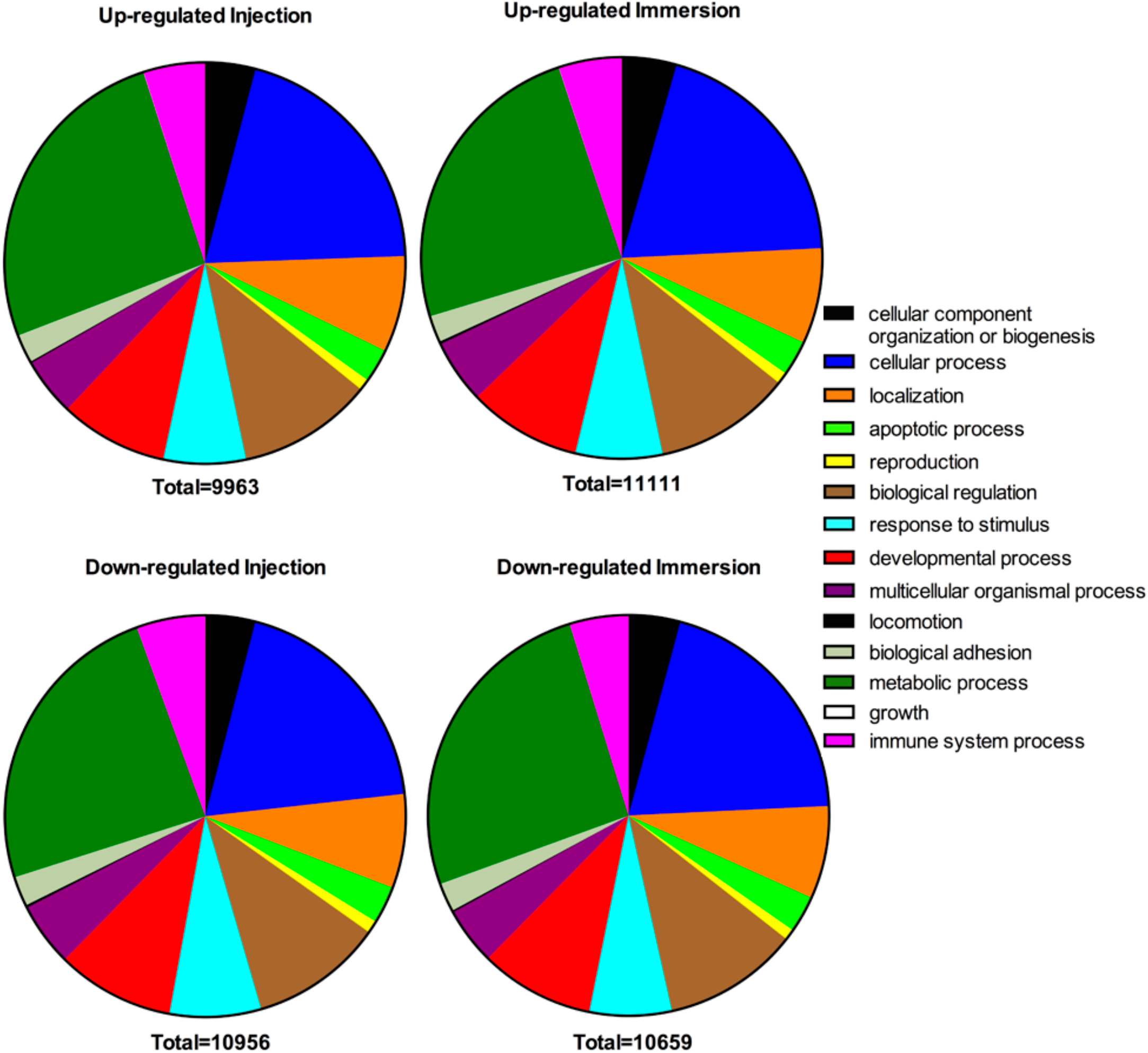
GO annotation of up-regulated and down-regulated proteins in zebrafish exposed to *P. aeruginosa* by injection or static immersion. Proteins significantly changed at 22 hpe in zebrafish exposed at 72 HPF by injection or static immersion with *P. aeruginosa* PAO1 were categorized by biological process (GO). The total represents the total matches in the database.

**Supplementary Figure 2.**
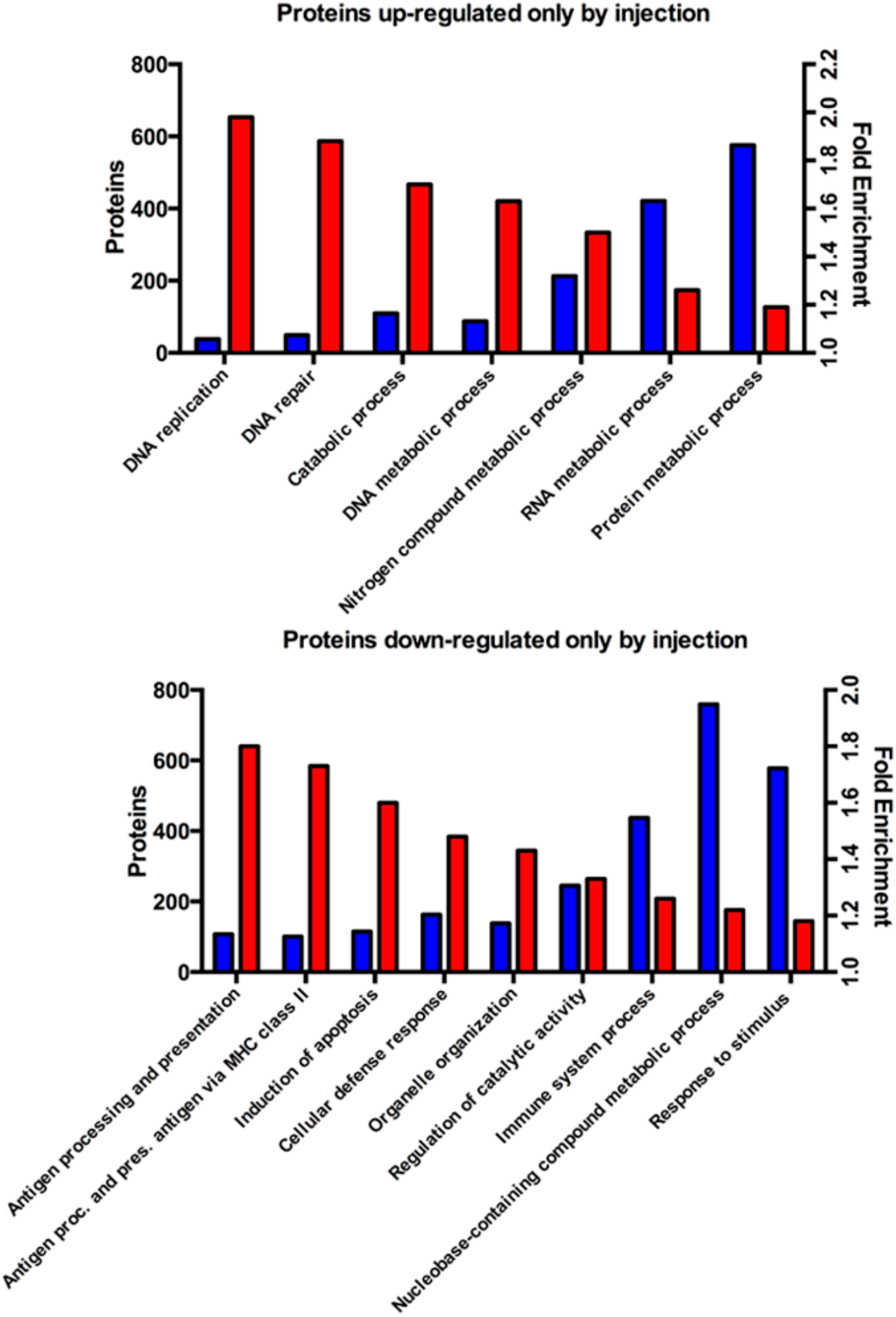
Overrepresentation analysis for proteins changed only in injection method. Up-regulated (top) or down-regulated (bottom) proteins at 22 HPI in zebrafish exposed at 72 HPF by injection (but not in larvae exposed by immersion) categorized by biological process (GO). Blue bars indicate number of proteins. Red bars indicate fold enrichment. The cut-off was set at P<0.05. Only the more relevant groups are presented. Proc. = processing. Pres. = presentation.

**Supplementary Figure 3.**
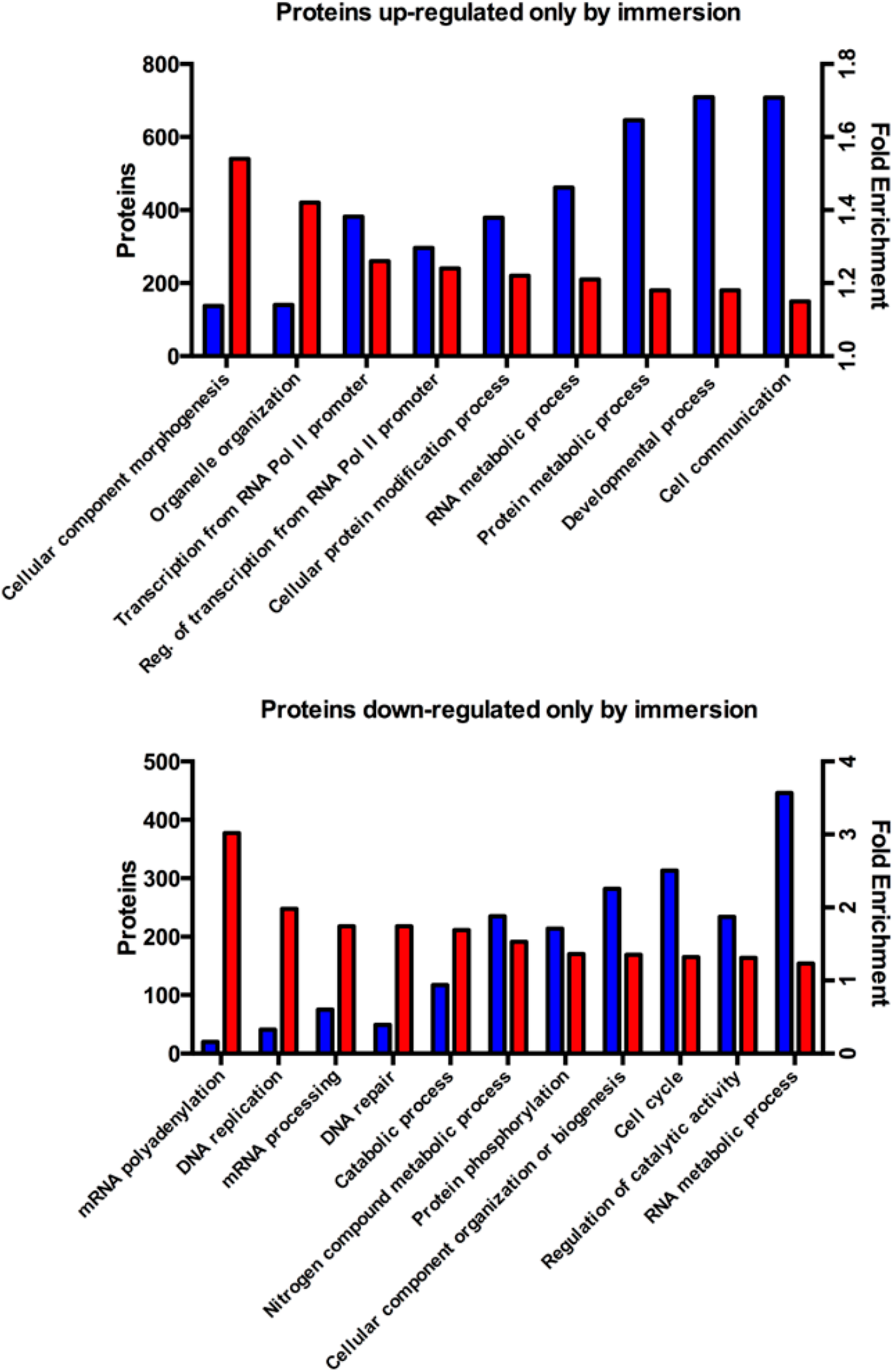
Overrepresentation analysis for proteins changed only in immersion method. Up-regulated (top) or down-regulated (bottom) proteins at 22 hpe in zebrafish exposed at 72 HPF by immersion (but not in larvae exposed by injection) categorized by biological process (GO). Blue bars indicate number of proteins. Red bars indicate fold enrichment. The cut-off was set at P<0.05. Only the more relevant groups are presented. Reg. = regulation.

